# Chimpanzees but not orangutans display aversive reactions toward their partner receiving a superior reward

**DOI:** 10.1101/274803

**Authors:** Yena Kim, Jae Choe, Gilsang Jeong, Dongsun Kim, Masaki Tomonaga

**Author notes:** Corresponding author: Yena Kim Corresponding author address: Research Institute of Ecoscience, Ewha Womans University, 52 Ewhayeodae-gil, Seodamun-gu, Seoul, 03760, Republic of Korea.

## Abstract

Fairness judgment is a fundamental aspect of human cooperation. By carefully balancing the payoffs and efforts with cooperating partner (s) we could either avoid or punish cheaters and stably maintain cooperation. Recent studies investigating the origin of this fairness sentiment have demonstrated that this psychological trait is not unique to humans, but also can be observed in other group-living primates, such as chimpanzees and capuchins, suggesting a convergent evolution of a sense of fairness, with cooperative social life being the selective pressure for it. The current study was designed to test this hypothesis by directly comparing the response to the outcome inequity in two of our closest living relatives, chimpanzees and orangutans, having different social systems, i.e. solitary and patrilocal multi-male multi-female groups. Unlike other inequity experiments, we used a prosocial choice apparatus with different reward distributions (advantageous / disadvantageous) to give subjects an active role of not-sharing foods if they considered it unfair. In addition to the choice, we also recorded the behavioral responses of the apes to the inequity. Throughout the experiments aversive emotional responses toward the disadvantageous inequity were only found in chimpanzees, but not in orangutans, supporting the convergent (or domain-specific) evolution of a sense of fairness. However, this aversion to the inequity did not lead the chimpanzees to actually make selfish choices, indirectly supporting the previous findings that chimpanzees employ a partner choice strategy rather than a punishment for fair cooperation. We also found that hierarchy seems to play an important role in the expression of aversion to inequity and prosocial tendency in chimpanzees.

## INTRODUCTION

Why humans cooperate and willingly suffer costs to punish free-riders has always been an evolutionary puzzle. Besides kin-selection (Hamilton 1964) and reciprocal altruism (Trivers 1971) to explain the ultimate cause of altruism, many attempts have been made to understand the proximate mechanisms of altruistic acts and altruistic punishment in human society using game-theoretic models. Social norms, one of the key features of human cooperation and altruism, have received particular attention (Fehr and Fischbacher 2004a, b). By enforcing social norms, cooperation can flourish and stabilize (Fehr and Fischbacher 2004a; Yamamoto and Takimoto 2012). Researchers have tried to understand what the psychological mechanisms are that enable and establish social norms in human society, and have suggested that fairness judgement modulating third party punishment plays a key role in this process (Fehr and Fischbacher 2004a, b). Altruistic social preference and fairness judgement are indeed mutually linked to each other, as demonstated in a series of games (e.g., dictator game, ultimatum game, and trust game, etc.; Lee 2008) and supported by neurophysiological studies. For example, mutual cooperation and altruistic punishment to social norm violators is known to be associated with brain regions responsible for reward processing (De Quervain et al. 2004; Rilling et al. 2002), and unfairness increases brain activity in the region involved in negative emotions (Sanfey et al. 2003). Developmental studies suggest that human infants as early as 12 months old are known to develop this sense of fairness, which suggests a genetically wired psychological system in human evolution (Geraci and Surian 2011). Based on this, we might wonder about the evolutionary roots of this sentiment of what is fair, to what extent we share this with other animals, and what are the physical or social conditions responsible for the evolution of fairness.

The first attempt to investigate the sense of fairness in other animals was made by Brosnan and de Waal using a token-exchange experiment with different quality foods given to monkeys as rewards (Brosnan and De Waal 2003). Since then, similar experiments have been replicated on multiple species of nonhuman primates, as well as other animal taxa (chimpanzees: Bräuer et al. 2006, 2009, Brosnan et al. 2005; capuchin monkeys: Dubreuil et al. 2006, Fletcher 2008, Roma et al. 2006, Silberberg et al. 2009; long-tailed macaques: Massen et al. 2012; squirrel monkeys: Talbot et al. 2011; dogs: Horowitz 2012, Range et al. 2009, Range et al. 2012; and ravens: Wascher and Bugnyar 2013). The results tend to support the link between species-specific sociality, such as their cooperative regime (species-specific sociality), and inequity aversion (Brosnan 2013). However, there still is an inconsistency within species across different study groups, possibly due to the different methodologies used in each study. For example, chimpanzees displayed sensitivity to unfair outcomes in a token-exchange experiment (Brosnan et al. 2005, 2010), as well as to unfair intentions in an experiment in which a choice was given to the chimpanzee subject to prevent sharing food with partners (Jensen et al. 2007b). However, another study using a modified ultimatum game to test inequity aversion in champanzees yielded negative results, demonstrating that chimpanzees do not refuse any unfair offers by a proposer unless it is a zero-food offer and therefore they are apparently not sensitive to unfair outcomes (Jensen et al. 2007a). There are many differences in the methodologies used, but the discrepancy was mainly driven by two important factors: 1) whether the task is intended to test outcome-based inequity or behavior-(or intention-) based inequity; and 2) whether the subject has a choice to prevent the partner’s free-riding. In the current study, we only address outcome inequity.

In the most widely tested token-exchange paradigm, two subjects sit side-by-side and perform an individual task in which they exchange tokens for a piece of food upon request by a human experimenter. In this setting, the subject receiving a reward inferior to that of her partner can either refuse to accept that reward or refuse to continue the task. In either case, no direct punishment (or active decision) by the subject towards the partner’s free-riding can be made. On the other hand, tasks using a food-delivering apparatus (Silk et al. 2005) allow subjects to make an active decision on whether or not to share or to divide rewards fairly with their partner. Therefore, in the latter setting, we can actually investigate whether the subject prevents unfair resource sharing, thereby punishing their partner’s free-riding. The additional advantage of the latter experimental setting is that the refusal or latency to do the task can also be measured in addition to the subject’s actual choice.

In the current study, we employed a food-delivering apparatus with differential reward distributions to test two things: 1) whether species-specific sociality plays an important role in the aversion to outcome inequity by comparaing two closely-related species of great apes, chimpanzees and orangutans, having different social systems; and 2) if aversion to inequity is observed, whether this aversion can be further expanded to the subject’s choice to prevent the partner from eating a superior reward without effort. We predicted that chimpanzees, a species which shows a variety of behavioral evidence for cooperation (Boesch 2003; Nishida 1983; Watts and Mitani 2002), but not orangutans, a species with a semi-solitary lifestyle (van Schaik 1999; van Schaik and van Hooff 1996;), would display aversive reactions toward outcome inequity, and thereby support the domain-specific evolution of aversion to inequity (Brosnan 2013).

## RESULTS

### Understanding of the task

To validate our experimental design and apparatus, we first looked at the apes’ understanding of operating the apparatus. Both chimpanzees and orangutans chose the double-rewarding option (or prosocial option when the partner was present: 1/1) over 90% of the time in the knowledge-test where there was no partner present, but access to the partner’s booth was allowed (chimpanzees: 91.95%, binomial test, *p* < 0.01; orangutans: 94.61%, binomial test, *p* < 0.01), suggeting that the apes understood the consequence of their choice delivering a food reward into the partner’s booth.

### Prosocial choices across differential reward distributions

Chimpanzees chose prosocial choices less often when they were paired with their partners than alone, regardless of reward distributions (GLMM; Table 1, Fig. 2), suggesting that chimpanzees behaved selfishly toward their partner in the prosocial-choice experiment. On the other hand, neither the reward distribution (advantageous/disadvantageous) nor the condition (partner present/partner absent) had an effect on orangutan prosocial choices (GLMM; Table 1, Fig. 2). When we looked at the individual level of prosocial choices by chimpanzees, we found an interesting difference in which Pendesa chose prosocial choices more often for her dominant partner (Puchi) than her subordinate partner (Mari), regardless of reward distributions (Mann-Whitney *U*-Test; N_mari_ = 8, N_puchi_ = 8, *U* = 12, *p* = 0.032; Fig. 3). This hierarchical influence was also consistently observed in Pendesa’s behavioral expressions in the disadvantageous reward distribution.

**Table 1:** Binomial GLMM results of prosocial choices of chimpanzees and orangutans in relation to condition (partner-present/partner-absent) and reward distribution (advantageous/disadvantageous)

| Prosocial choices |  |  |  |  |  |
| --- | --- | --- | --- | --- | --- |
| Species | Predictor variable |  |  |  |  |
| Chimpanzees | | Estimate | SE | Z | $P(>\vert z\vert )$ |
|  | (Intercept) | 0.1772 | 0.1923 | 0.921 | 0.357 |
|  | Condition<br>(partner present) | -0.4155 | 0.2072 | -2.005 | 0.045* |
|  | Reward distribution<br>(disadvantageous) | 0.3502 | 0.2072 | 1.690 | 0.091 |
| Orangutans | | Estimate | SE | Z | $P(>\vert z\vert )$ |
|  | (Intercept) | 0.0038 | 0.2306 | 0.017 | 0.986 |
|  | Condition<br>(partner present) | 0.1352 | 0.2644 | 0.511 | 0.609 |
|  | Reward distribution<br>(disadvantageous) | 0.4372 | 0.2645 | 1.653 | 0.098 |

**Fig. 2:**
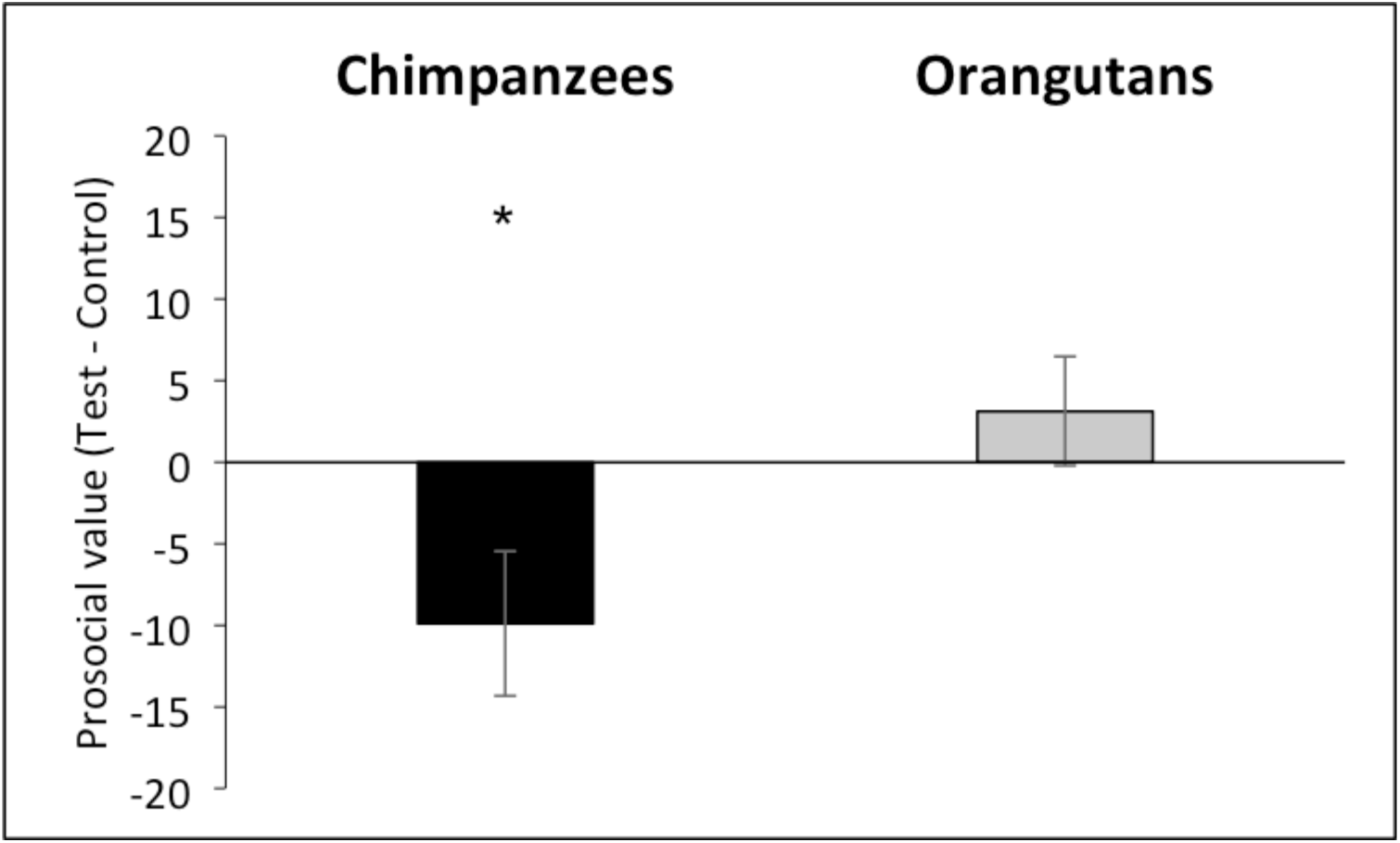
Prosocial value (±SE) for chimpanzees and orangutans. Mean percentage of prosocial choices in the control-test (partner-absent condition) subtracted from the mean percentage of prosocial choices in the prosociality-test (partner-present condition) regardless of reward distributions. Positive value indicates higher prosocial choices in the partner-present condition than in the partner-absent condition. *: *p* < 0.05.

**Fig. 3:**
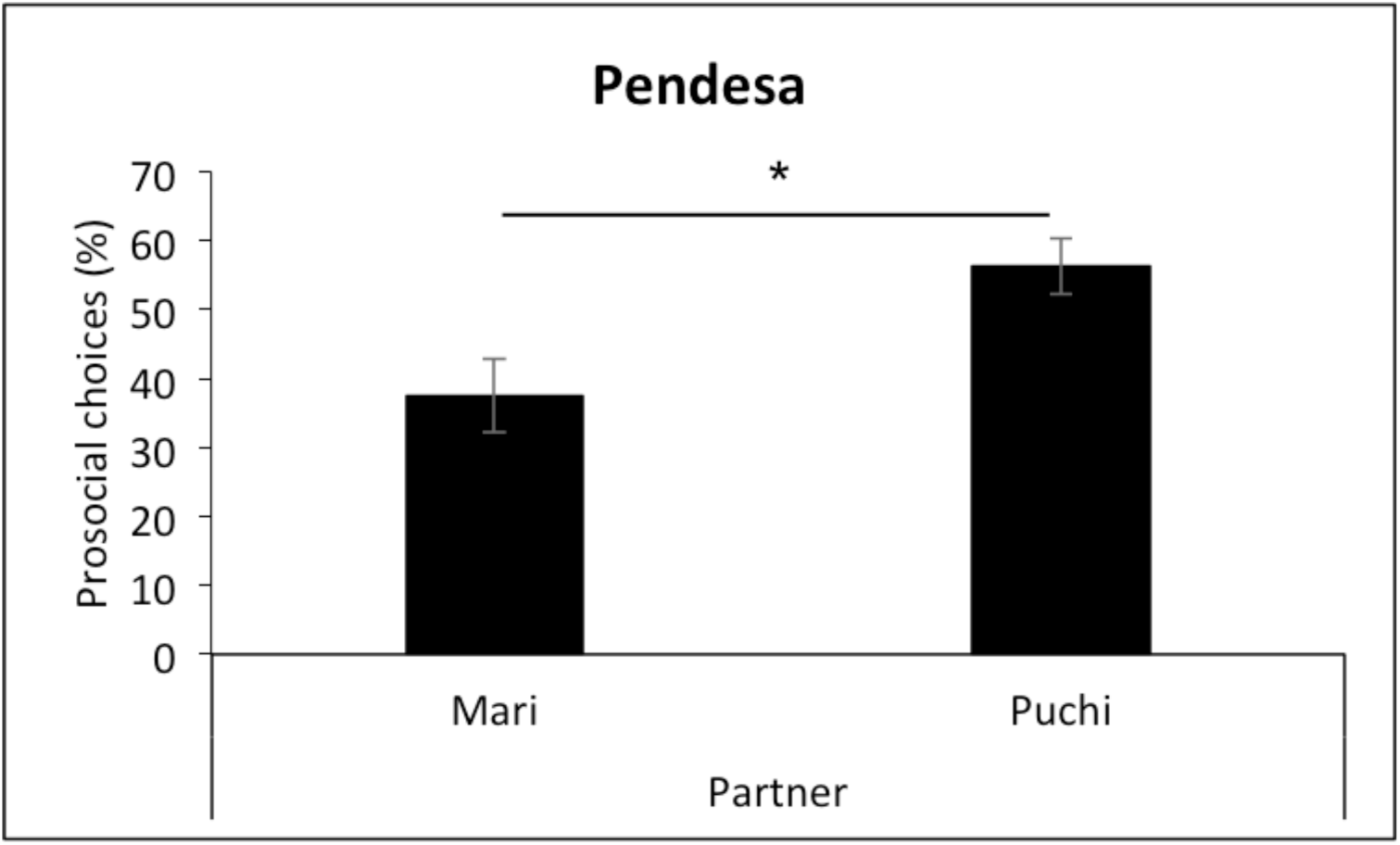
Mean percentage of prosocial choices (±SE) in a session by Pendesa in the prosociality-test (partner-present condition) with different partners regardless of the reward distribution. *: *p* < 0.05.

### Latency across differential reward distributions

Both chimpanzees and orangutans in general took longer to complete a trial when they were in a disadvantageous reward distribution and also when they were paired with their partners, but we found no interaction (GLMM; Table 2, Fig. 4a, b), perhaps indicating that less-preferred rewards in general elicited the actor’s aversion to the task. A partner’s influence on the longer latency as well could be simply interpreted as an increase in the actor’s attention or interaction directed to the partner, instead of aversion to the partner’s free-riding.

**Table 2:** Negative binomial GLMM results of latency to complete a trial by chimpanzees and orangutans in relation to condition (partner-present/partner-absent) and reward distribution (advantageous/disadvantageous).

| Latency to complete a trial |  |  |  |  |  |
| --- | --- | --- | --- | --- | --- |
| Species | Predictor variable | Estimate | SE | Z | $P(>\vert z\vert )$ |
| Chimpanzees | (Intercept) | -0.3050 | 0.2435 | -1.253 | 0.2103 |
|  | Condition (partner-present) | 0.3166 | 0.1344 | 2.356 | 0.0185* |
|  | Reward distribution (disadvantageous) | 1.5777 | 0.1400 | 11.269 | <2e-16*** |
| Orangutans | (Intercept) | 0.1578 | 0.2069 | 0.763 | 0.446 |
|  | Condition (partner-present) | 0.5431 | 0.1395 | 3.893 | 9.91e-05*** |
|  | Reward distribution (disadvantageous) | 1.0228 | 0.1452 | 7.043 | 1.88e-12*** |

**Fig. 4:**
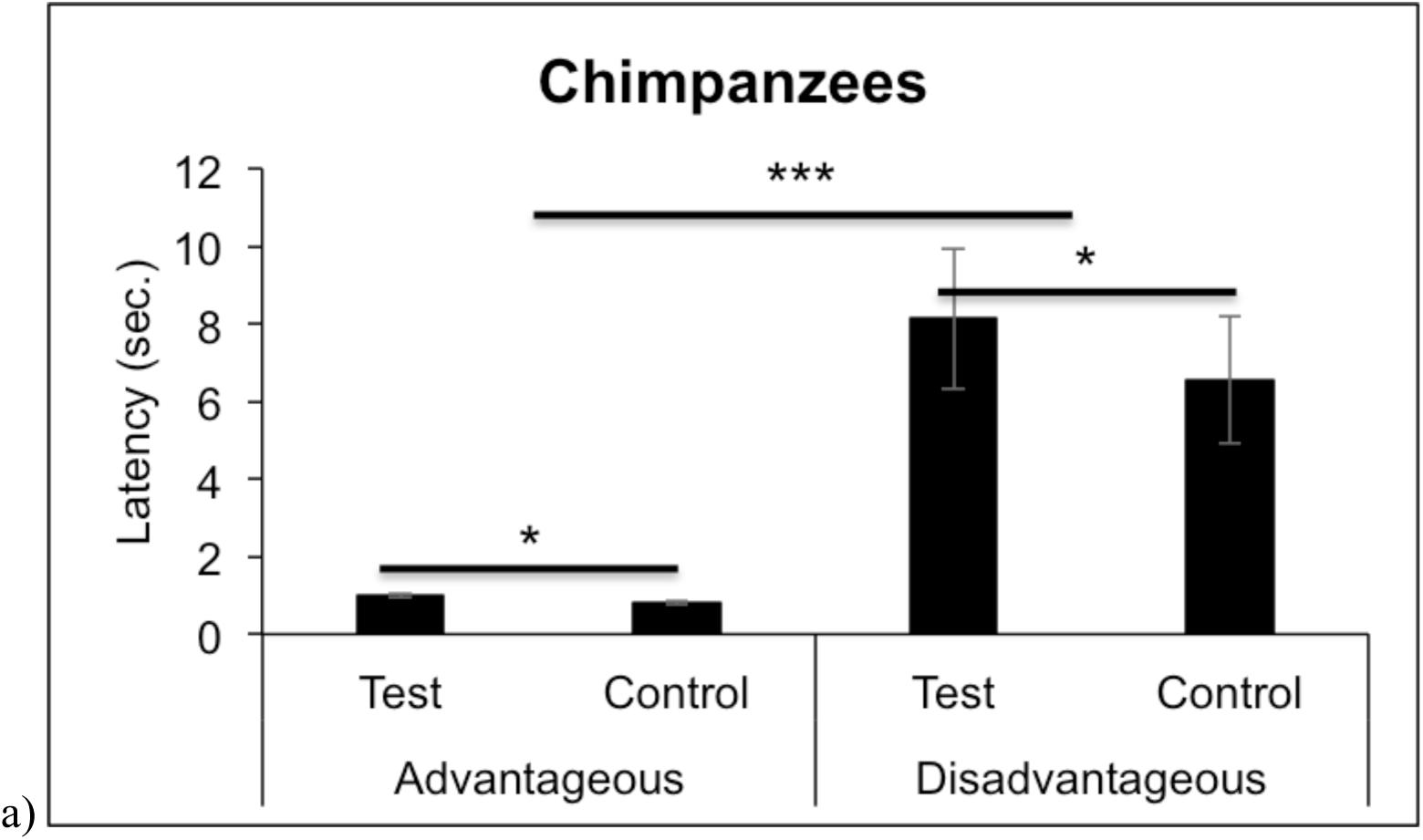
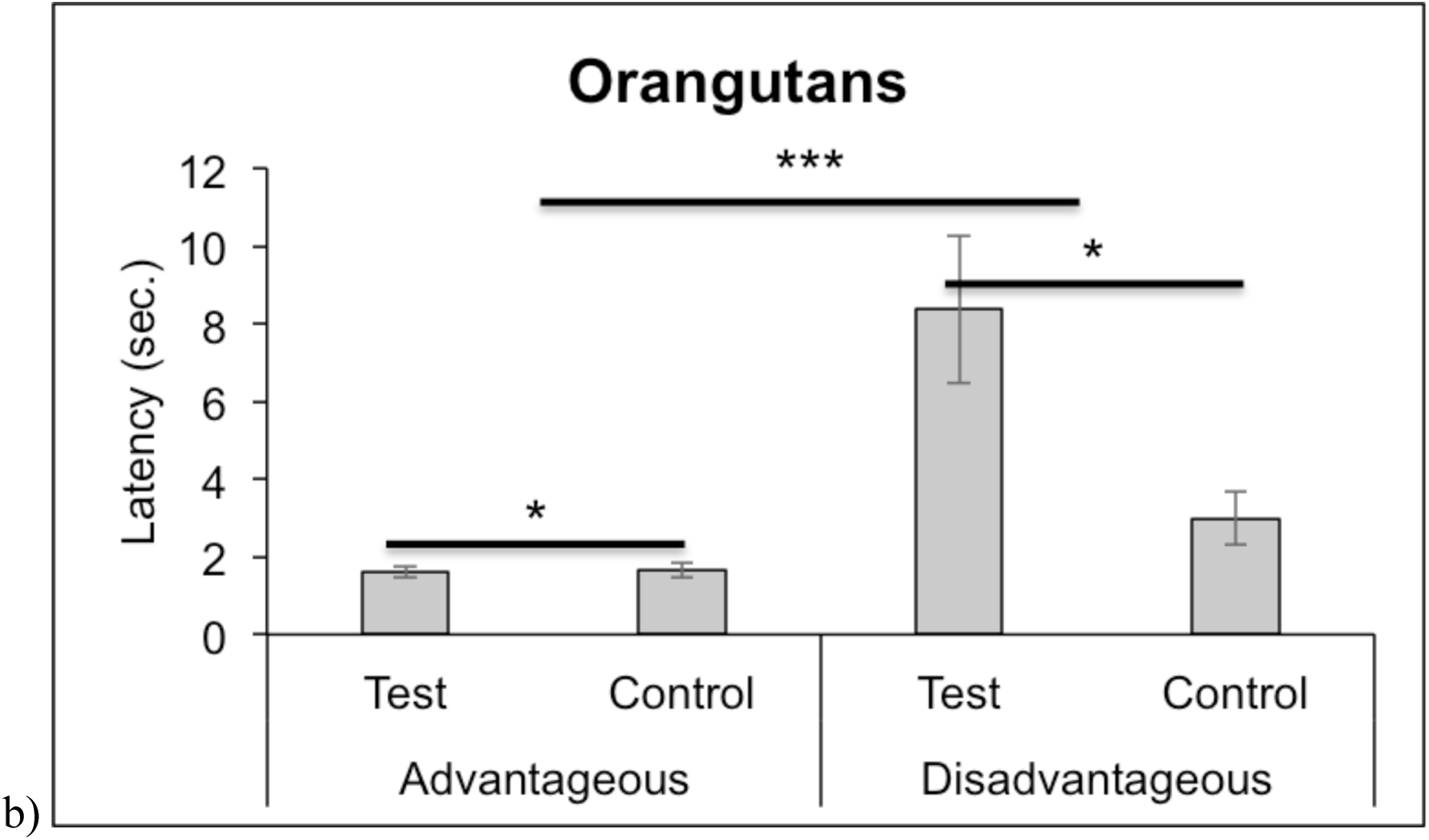
Mean latency (±SE) to complete a trial by chimpanzees (a) and orangutans (b) in relation to condition (partner present/partner absent) and reward distribution (advantageous/disadvantageous). *: *p* < 0.05.

### Aggressive display across differential reward distributions

In addition to latency, we also recorded the actor’s displaying any sign of aggressive behavior directed to its partner. We found only one pair of chimipanzees, Pendesa–Mari, where Pendesa was dominant over Mari, diplaying aversive reactions in the disadvantageous reward distribution. It is important to note that: 1) Pendesa never displayed to Mari in the advantageous reward distribution, and 2) Pendesa was not observed to display in the partner-absent (control) condition. Moreover, when she was paired with a dominant partner (Puchi), her displaying diappeared completely in all conditions. The total number of aggressive displays directed by Pendesa to Mari was 13 in 32 disadvantageous trials (34.43%; 4 sessions of the 8-trial prosociality test), and more specifically, Pendesa did not display at all when she was making selfish choices toward Mari (preventing Mari’s free-riding), but displayed in almost all cases (84.61%; 11 times out of 13 prosocial choices) when she was making prosocial choices toward Mari.

## DISCUSSION

In the current study, we investigated the species difference in aversion to outcome inequity. Unlike other inequity experiments using a token-exchange paradigm, we did not design our experiment to test for an effect of effort on the aversion to inequity, in addition to the outcome disparity. Rather, we manipulated the outcome in only two opposing ways, with an 1) advantageous reward distribution versus a 2) disadvantageous reward distribution for the actor, thereby maximizing the contrast between these two conditions. Moreover, by requiring only the actor to make an effort to operate the apparatus, together with the disadvantageous reward distribution, we expected to observe a higer likelihood of expressing aversion to inequity.

By introducing the knowledge test, we validated the apes’ understanding about our procedure and the apparatus, which was demonstrated by a higher number of double-rewarding choices (or prosocial choices when the partner was present: 1/1) by the chimpanzees and the orangutans in the knowledge-test. If the subjects are averse to the disadvantageous outcome inequity strong enough to prevent the partner’s free-riding, we would expect the actors to choose selfish choices more frequently in the condition where food rewards are disadvantageously distributed, especially when the parnter is present than when absent. In the same manner, we would also expect that longer latency would be observed in the disadvantageous reward distribution with the partner present.

However, we found no evidence that the disadvantageous reward distribution elicited selfish choices in either chimpanzees or orangutans.

There was a difference in terms of overall prosocial choices between the two species, in which chimpanzees made fewer prosocial choices in the partner-present (prosociality-test) condition than the partner-absent (control) condition, but no difference was observed in orangutans, suggesting perhaps a general aversion to the partner’s free-riding on food rewards in chimpanzees. This could in part explain the negative results in experiments involving a prosocial-choice apparatus (Silk et al. 2005; Vonk et al. 2008; Yamamoto and Tanaka 2009, 2010), as opposed to the observational evidence of chimpanzee prosociality (Boesch 1994; Watts 2000; Wittig et al. 2014) or instrumental helping in experimental settings (Warneken and Tomasello 2006; Yamamoto et al. 2009, 2012). Indeed, chimpanzees are known to divide the carcass obtained during collective hunting, based on each individual’s contribution to the hunt, though hierarchy or social relationships also influence carcass distribution (Watts and Mitani, 2002). Chimpanzees are also able to select collaborating partners who have been helpful before, suggesting their ability to remember and recognize a partner’s previous contribution (Melis et al, 2006). Therefore, it might not be so surprising to see the chimpanzees in our experiment behave selfishly toward the partners who did not contribute earlier to their food acquisition.

One might argue that the chimpanzee’s higer prosocial choices in the partner-absent condition than in the partner-present condition resulted from chimpanzees’ expectation of taking the food reward from the partner’s booth after the experiment. However, if that was the case, we could expect this behavior to diminish over the progression of the sessions, since they would learn there is no way they can get the rewards from the partner’s booth. However, we found no difference in prosocial choices between the first and the last sessions of the partner-absent (control) conditions (Mann-Whitney *U*-test, *U* = 16, *P* = 0.4134).

We also checked latency to complete a trial to see whether the chimpanzees and orangutans in our experimental setup expressed any sign of aversion to outcome inequity. Interestingly, both species showed a similar pattern of longer latency in the partner-present (prosociality-test) condition than partner-absent (control-test) condition, and disadvantageous than advantageous reward distribution. However, the absence of interaction between these two factors in our model makes it difficult to conclude that the longer latency observed in the disadvantageous reward distribution was caused by the partner getting a more superior reward than the actor, instead of a general aversion to the less-preferred rewards or possibly a frustration effect (Dubreuil et al, 2006). Increased latency in the partner-present condition, compared to the partner-absent condition, also could be explained otherwise; i.e. increased attention or interactions with partners.

However, we found a pair of chimpanzees, Pendesa–Mari, where Pendesa was dominant over Mari, showing a clear sign of aversion to inequity caused by the partner getting a more superior reward than the actor. Pendesa’s display was not observed in any of the conditions where the foods were advantageously distributed, when the partner was absent, or when Puchi, who was dominant over Pendesa, was paired with her, suggesting her reaction is clearly partner-and reward-dependent. Especially, her displays were only observed when Mari was taking a food reward, thanks to Pendesa’s prosocial choices, but never in selfish choices. This interesting finding was also supported by Pendesa’s overall choices in which Pendesa made the selfish choice more for her subordinate partner, Mari, than her dominant partner, Puchi. Moreover, the consistent behavioral tendency observed in the same Pendesa–Mari pair with a different test paradigm, a tool-transfer experiment (Yamamoto et al. 2009), suggests a consistent social influence on prosociality across time, regardless of task paradigms (Brosnan et al. 2005; Massen et al. 2012; Cronin 2012). The fact that we did not find any sign of aggression in the Chloe–Cleo pair could be attributed to their kin relationship.

To bear this in mind, our results, as predicted by Brosnan’s hypothesis of convergent (domain specific) evolution of inequity aversion (Brosnan 2013), suggest that chimpanzees, a species having complex sociality with a cooperative regime, have a sense of fairness, that orangutans do not seem to possess, though whether orangutans have lost or chimpanzees have acquired this trait in the hominid lineage remains to be tested together with the studies in gorillas as well. More specifically, chimpanzees may exhibit emotional responses towards their partner who is taking superior reward than them with no effort, but this does not lead them to make selfish choices. Furthermore, the general selfish tendency observed in these chimpanzees toward their partners in the prosocial-choice experiment suggests that a prosocial tendency in chimpanzees can be masked by the imbalance of effort among individuals involving cooperative or prosocial behaviors. Even though our experiment did not test third-party punishment directly (but see Riedl et al, 2012), the fact that chimpanzees did not show an active punishment in dyadic relationships when it comes to outcome disparity, clearly distinguish us from chimpanzees. Further research should be done to test whether the sensitivity to outcome inequity observed in chimpanzees can be further generalized to behavioral (intention-based) inequity, and to understand more specifically about the nature of the sense of fairness and aspects of social influence.

## METHODS

### Study site and participants

Three pairs of chimpanzees (Chloe–Cleo, Pendesa–Puchi, Pendesa–Mari; 5 females) from the Primate Research Institute (PRI, hereafter), Kyoto University, Japan participated in this experiment. All individuals housed at PRI have voluntary access to enriched indoor and outdoor environments (Matsuzawa 2006; Ochiai and Matsuzawa 1997). Chloe (GAIN ID: 0441) and Cleo (GAIN ID: 0609) form a mother-offspring pair and Pendesa (GAIN ID: 0095), Puchi (GAIN ID: 0436) and Mari (GAIN ID: 0274) are from the same social group (綿貫宏史朗 et al. 2014). Pendesa was dominant over Mari but subordinate to Puchi during the study period according to the caretakers (Yamamoto et al. 2009). All participants had previously participated in a variety of cognitive experiments using computerized touch panels, but had never been exposed to this prosocial-choice experiment using a food-delivering apparatus. The study was carried out from December 2014 to March 2015, before the chimpanzees’ mid-day feeding. They were not water or food deprived.

Two pairs of orangutans (Bosuk–Bora, Bora–Boram; 2 males: Bosuk and Boram,1 female: Bora) from the Seoul Zoo, Republic of Korea, also participated in the experiment. Bosuk and Bora had been housed together for more than 10 years and Boram was housed alone due to aggression received from adult male Bosuk. All individuals previously participated in a prosocial-choice experiment using the same apparatus, but without reward manipulations (Kim et al. 2015). The study was carried out from August to October 2015 before the orangutans were released into the enriched outdoor area for the exhibition. They were not water or food deprived.

### Experimental setting and apparatus

The chimpanzee and orangutan participants were tested in two adjecent booths (W x L x H: 120 x 260 x 250 cm and 170 x 260 x 250 for PRI, and 110 x 140 x 200 cm and 165 x 140 x 200 cm for Seoul Zoo) with either a plexiglass wall (PRI) or meshed wall (Seoul Zoo) in-between, which allowed the participants to visually and vocally communicate with each other. Only one of the pair, the actor, was given a choice to operate the apparatus while the other individual, the partner, sat passively next to the actor waiting for the choice to be made. We employed a food-delivering apparatus (or prosocial-choice apparatus) which was previously used to test voluntary food sharing in captive orantuans (Kim et al. 2015). The box-shaped appratus, made of steel bars (W x L x H: 100 x 30 x 65 for PRI and 120 x 30 x 65 cm for Seoul Zoo) with 4 wheels on the bottom was positioned in front of the participants seated in the two booths. In the front (facing the cage) of the apparatus, there were four square food holes (5 x 5 cm) in a line, two in one half of the apparatus allocated for the actor and the other two on the other half of the apparatus allocated for the partner. One big transparant plexiglass (W x L: 121 x 21 for PRI and 140 x 22 cm for Seoul Zoo) wall with two pierced rectangular holes (W x L: 18 x 5 cm; windows, hearafter), one positioned in the middle of the two food holes for the actor and another for the partner, was attached to the front of the apparatus. A handle under the window allowed the actor to slide open the windows, either to the left or right side of the food holes to retreive the food rewards inside. Either of the two food holes in the partner side could be covered by opaque occluders, giving the partner only one side option to access the food reward. The position of the occluders was randomized and counterbalanced across trials.

### Procedures

One experimental session consisted of three different series of tests in succession: 8 trials of the knowledge-test, 8 trials of the prosociality-test, and 8 trials of the control-test. Each pair underwent one experimental session per day and had 4 sessions in total for each reward distribution (advantageous/disadvantageous). Advantageous reward distribution (Fig. 1a) is where two preferred food rewards (two less-preferred food rewards in the disadvantageous reward distribution) are baited in the two food holes for the actor and one less-preferred reward and one occluder (one preferred food reward and one occluder in the disadvantageous reward distribution; Fig. 1b) are baited in each of the food holes for the partner. Each pair had 4 sessions of 8-trial advantageous reward distribution and 4 sessions of 8-trial disadvantageous reward distribution. The order of the sessions was randomized and counterbalanced. We used the same apparatus to determine food preferences of the actor by giving a dichotomous choice between different food items in the food holes. Each participant received 10 trials with two food items and a food item was chosen as preferred, if the participant chose over 90% of one item. Chimpanzees preferred a peanut over a piece of carrot and orangutans preferred a piece of dried pineapple over a peanut. We first tested orangutans with a piece of sweet potato and a peanut which were included in the orangutans’ daily diet, but they did not eat sweet potatos at all in the experimental room and therefore we decided to test with peanuts and dried pineapples.

**Fig. 1:**
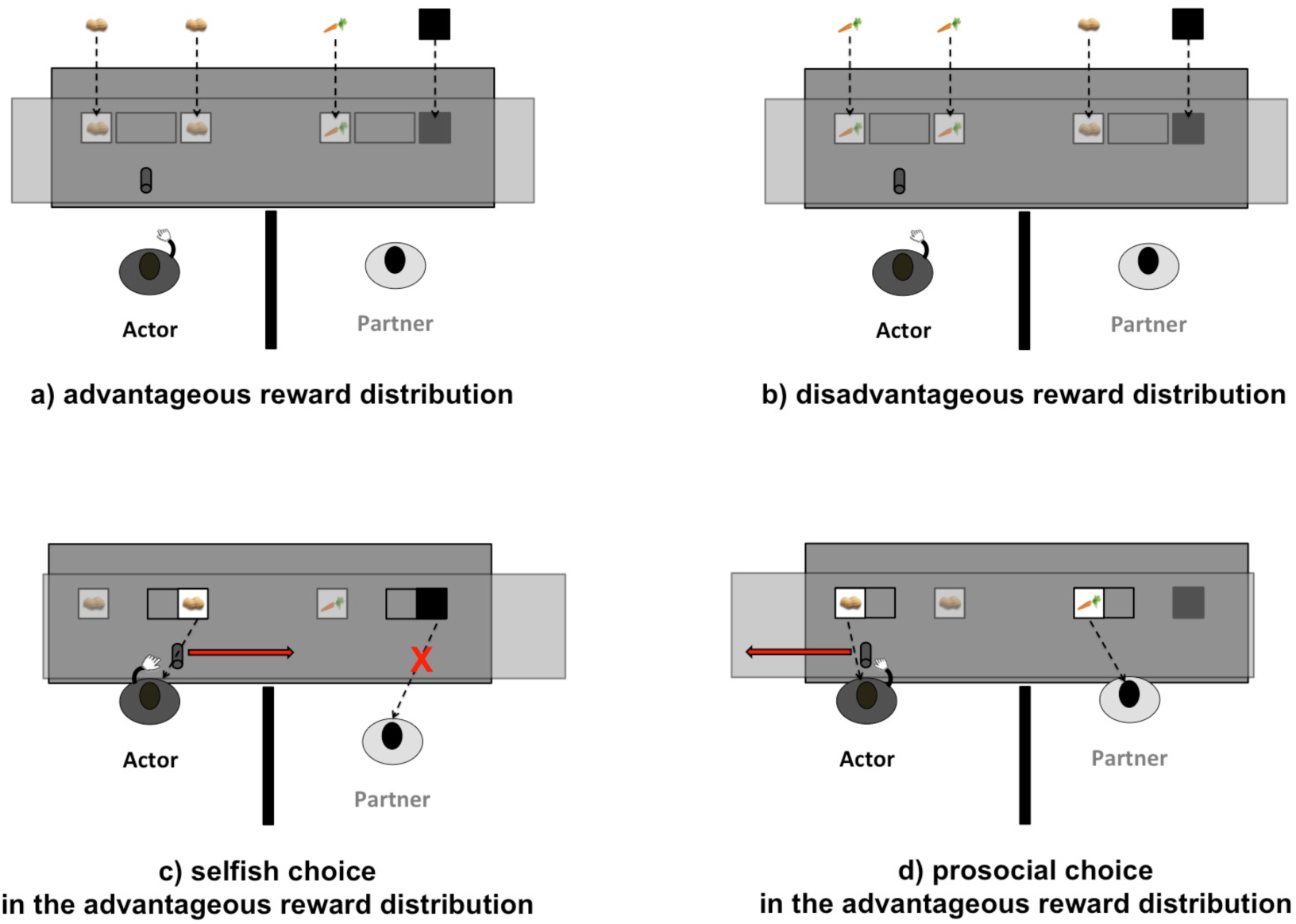
Illustration of the experimental setting with differential reward distributions and operation of the apparatus. a) Advantageous reward distribution: two preferred rewards (peanuts in this figure) were baited for the actor and one less-preferred reward (a carrot in this figure) and an occluder were baited for the partner. b) Disadvantageous reward distribution: two less preferred rewards were baited for the actor and one preferred and an occluder were baited for the partner, when the less-preferred reward for the partner was located on the left side of the food holes and the occluder was on the right side of the holes. c) Demonstrates a selfish choice where the actor slides the door to the right (occluder side). d) Demonstrates a prosocial choice where the actor slides the door to the left (food side)

### Knowledge-test

In the knowledge test, only the actor was given the apparatus without a partner, with access to the partner-side also. Two less-preferred food rewards were baited in the actor-side food holes, and one preferred food reward and one occluder in the partner-side food holes. If the actor chose a double-rewarding choice (1/1: one food reward from the actor-side and another food reward from the partner-side food holes), this indicates that the actor understood the consequence of the choice and how to operate the apparatus. The experimental session was started with 8 trials of knowledge-tests, regardless of the reward distribution, to make sure the actors in every session understood the task. The criterion to proceed to the test phase was 7 double-rewarding choices out of 8 trials. If they missed more than 2 trials, we added two trials more and if they missed more than that, we did not continue with the experimental session on that day.

### Prosociality-test (partner-present condition)

In the prosociality-test, the door between the two booths was closed and the partner allowed to enter the booth next to the actor. Only the actor was able to make either a selfish choice (1/0: only one food reward was available for the actor and no food with only an occluder available for the recipient) or a prosocial choice (1/1: double-rewarding choice in the knowledge test).

### Control-test (partner-absent condition)

In the control-test, the door between the two booths remained closed but the partner was absent. Therefore, the prosocial choice (1/1) by the actor did not provide any benefits to the actor or partner. We compared the prosociality and control test to measure the prosocial tendency of the actor.

## Coding and analysis

All experimental sessions were videotaped using two video cameras (Sony HDR-CX560), one from the front and another from the back. The choice (1/1 vs. 1/0) made by the actor was recorded by the research assistant as it ocurred. Latency is commonly used as a variable to test the aversion to inequity in token-exchange expeiments. In this experiment, we also measured latency to complete a trial as an indirect measure of aversion to inequity and tested whether the reward distribution and condition had an effect on the latency for each species. We also recorded the behavioral expression (aggressive display directed to the partner) by the actor. A binomial test was performed to examine the subject’s understanding of the task. For the analysis of prosocial choice data, we performed a GLMM for binary responses for each species including reward distribution (advantageous/disadvantageous), condition (partner present: prosociality test/partner absent: control test), and their interaction as predictor variables, and session number and pair as random effects. For the latency data, we also performed a GLMM fitted with a negative binomial error distribution for each species. The model included reward distribution, condition, and their interaction as predictor variables, and session number and pair as random effects. We compared the full models of prosocial choice and latency with the null models without predictor variables but containing random effects to assess the significance of the predictor variables, using a likelihood ratio test (*p* < 0.05 for both models). We also performed a Mann-Whitney *U*-test to compare prosocial choices between the pairs Pendesa–Mari and Pendesa–Puchi, and between the first and the last sessions of partner-absent (control) conditions. All *p-values* reported are two-tailed (α = 0.05), except the binomial test for the knowledge-test (one-tailed)since we predict that the subjects would choose higher double-rewarding choices than the chance level (50%).

## ACKNOWLEDGEMENTS

This research was financially supported by KAKENHI grant (#23220006, 15H05709 to MT and #25666 to YK), the Ewha Global Top5 Grant 2013 of Ewha Womans University, and Amore Pacific Academic and Cultural Foundation, Korea. We thank the caretakers at the Seoul Zoo, Yangmuk Lim, Dongsun Kim, Jonghwa Lee, and Yoo Rok Park, and Primate Research Institute, Kyoto University, Norihiko Maeda, Shizuka Godjari, Yui Fujimori, and Atsushi Yamanaka, and the primate team of Ewha Womans University, especially Yoonjung Yi and Ahyun Choi, for their assistance. We are also grateful to Dr. Michael A Huffman for his invaluable comments on the manuscript and Dr. Heungjin Ryu and Andrew McIntosh for their help with the statistics. This study was approved by the Animal Welfare and Animal Care Committee of the Primate Research Institute, Kyoto University and by the Seoul Zoo Institutional Animal Care and Use Committee (IACUC no. 2016-001).

